# Brain activity reveals multiple motor-learning mechanisms in a real-world task

**DOI:** 10.1101/2020.03.04.976951

**Authors:** Shlomi Haar, A. Aldo Faisal

**Affiliations:** Brain and Behaviour Lab: Dept. of Bioengineering, Imperial College London, London, UK; Dept. of Computing, Imperial College London, London, UK; UKRI CDT in AI for Healthcare, Imperial College London, London, UK; MRC London Institute of Medical Sciences, London, UK

**Keywords:** motor learning, skill, real-world, EEG, post-movement Beta rebound, motor neuroscience

## Abstract

Many recent studies found signatures of motor learning in neural Beta oscillations (13– 30Hz), and specifically in the post-movement Beta rebound (PMBR). All these studies were in controlled laboratory-tasks in which the task designed to induce the studied learning mechanism. Interestingly, these studies reported opposing dynamics of the PMBR magnitude over learning for the error-based and reward-based tasks (increase versus decrease, respectively). Here we explored the PMBR dynamics during real-world motor-skill-learning in a billiards task using mobile-brain-imaging. Our EEG recordings highlight the opposing dynamics of PMBR magnitudes (increase versus decrease) between different subjects performing the same task. The groups of subjects, defined by their neural dynamics, also showed behavioural differences expected for different learning mechanisms. Our results suggest that when faced with the complexity of the real-world different subjects might use different learning mechanisms for the same complex task. We speculate that all subjects combine multi-modal mechanisms of learning, but different subjects have different predominant learning mechanisms.

## Introduction

Many different forms of motor learning were described and studied using various laboratory-tasks over the past decades (for review see Krakauer et al., 2019). Two main learning mechanisms are considered to account for most of our motor learning capabilities: error-based adaptation and reward-based reinforcement learning. Error-based adaptation is driven by sensory-prediction errors, while reward-based learning is driven by reinforcement of successful actions (Krakauer and Mazzoni, 2011). While both mechanisms can contribute to learning in any given task, the constraints of the highly controlled laboratory-tasks common in the field induce the predominance of one mechanism over the other (Haith and Krakauer, 2013), and show different neural dynamics associated with the different learning mechanisms (e.g. Uehara et al., 2018; Palidis et al., 2019).

The main neural signatures of voluntary movement and motor learning found in constrained laboratory tasks are the Beta oscillations (13–30 Hz), which are related to GABAergic neural activity (Roopun et al., 2006; Yamawaki et al., 2008; Hall et al., 2010, 2011). More specifically, there is a transient and prominent increase in Beta oscillations magnitude across the sensorimotor network after cessation of voluntary movement known as post-movement Beta rebound (PMBR) or post-movement Beta synchronization (Pfurtscheller et al., 1996). In motor adaptation studies, PMBR over the motor cortex contralateral to the moving hand was reported to negatively correlate with movement errors, lower errors induced higher PMBR (e.g. Tan et al., 2014a, 2016; Torrecillos et al., 2015) and therefore PMBR increases over learning. In reward-based tasks the PMBR shows the opposite trend; e.g., in a force tracking task PMBR decreased with learning (Kranczioch et al., 2008). Additionally, PMBR is positively correlated with GABA concentration as measured by magnetic resonance spectroscopy (Gaetz et al., 2011; Cheng et al., 2017) which also decreases over reward-based learning tasks such as sequence learning in force tracking (Floyer-Lea et al., 2006) and serial reaction time (Kolasinski et al., 2019).

We are now seeking to understand to what extent previous findings in artificial laboratory-tasks can be validated in a complex, fully-body task people choose to experience in daily life. Here, we set to study the human brain activity during motor learning in a real-world task using mobile EEG, We recently introduced a real-world motor-skill learning paradigm in pool table billiards (Haar et al., 2019). Here, we set to study the human brain activity during motor learning in a real-world task using mobile EEG. Subjects had to do a pool shot to put the ball in the pocket using full-body, self-paced movement, with as many preparatory movements as the subject needs for each shot. We implemented this as a real-world task because we are basically only adding sensors to a pool table setting. Subjects use the natural tools and setups they normally would, carry out the natural motor commands, receive the natural somatosensory feedback and experience the same satisfaction rewards when they put the ball in the pocket. In our pool playing paradigm, as in most everyday motor learning experiences, performance errors were not driven by artificial perturbations but by the complexity of learning the task (which takes competitive pool players years to master) and noise in the nervous system (Faisal et al., 2008). We test here the hypothesis whether neural correlates of motor learning in real-world tasks show features consistent with those in artificial laboratory tasks. Specifically, we hypothesize that PMBR responses may look different in real-world tasks, because learning in a real-world paradigm may not be predominantly mediated by a single specific learning mechanism, such as motor adaptation and its increasing PMBR response over trials. Moreover, we hypothesise that a far less constrained real-world task, may give human subjects the freedom to learn in their personally most conducive way, instead of being forced by an artificial paradigm to explore a single route of leaning: thus we want to test the hypothesis if different subjects may employ different learning strategies and consequently exhibit different neural signatures of learning or if all learn the same way.

## Methods

### Experimental Setup and Design

30 right-handed healthy human volunteers (12 women and 18 men, aged 24±3) with normal or corrected-to-normal visual acuity participated in the study. The recruitment criteria were that they played pool/billiards/snooker for leisure fewer than 5 times in their life, never in the recent 6 months, and had never received any pool game instructions. All volunteers gave informed consent before participating in the study, and all experimental procedures were approved by the Imperial College Research Ethics Committee and performed in accordance with the declaration of Helsinki. The volunteers stood in front of a 5ft pool table (Riley Leisure, Bristol, UK) with 1 7/8” (48mm diameter) pool balls. Volunteers performed 300 repeated trials where the cue ball (white) and the target ball (red) were placed in the same locations. We asked volunteers to shoot the target ball towards the pocket of the far-left corner (Figure 1A). Trials were split into 6 sets of 50 trials with a short break in-between to allow the subjects to rest a bit and reduce potential fatigue. Each experimental set (of 50 trials) took 8 to 12 minutes. For the data analysis, we further split each set into two blocks of 25 trials each, resulting in 12 blocks. During the entire learning process, we recorded the subjects’ brain activity with a wireless EEG headset (Figure 1B). The balls on the pool table were tracked with a high-speed camera to assess the subjects’ success in the game and to analyse the changes throughout learning, not only in the body movement and brain activity but also in its outcome – the ball movement (Figure 1C). EEG and ball motion tracking camera were recorded on the same machine. All signals were time-stamped by accessing the high precision event timer of the computer and synchronised accordingly.

**Figure 1.**
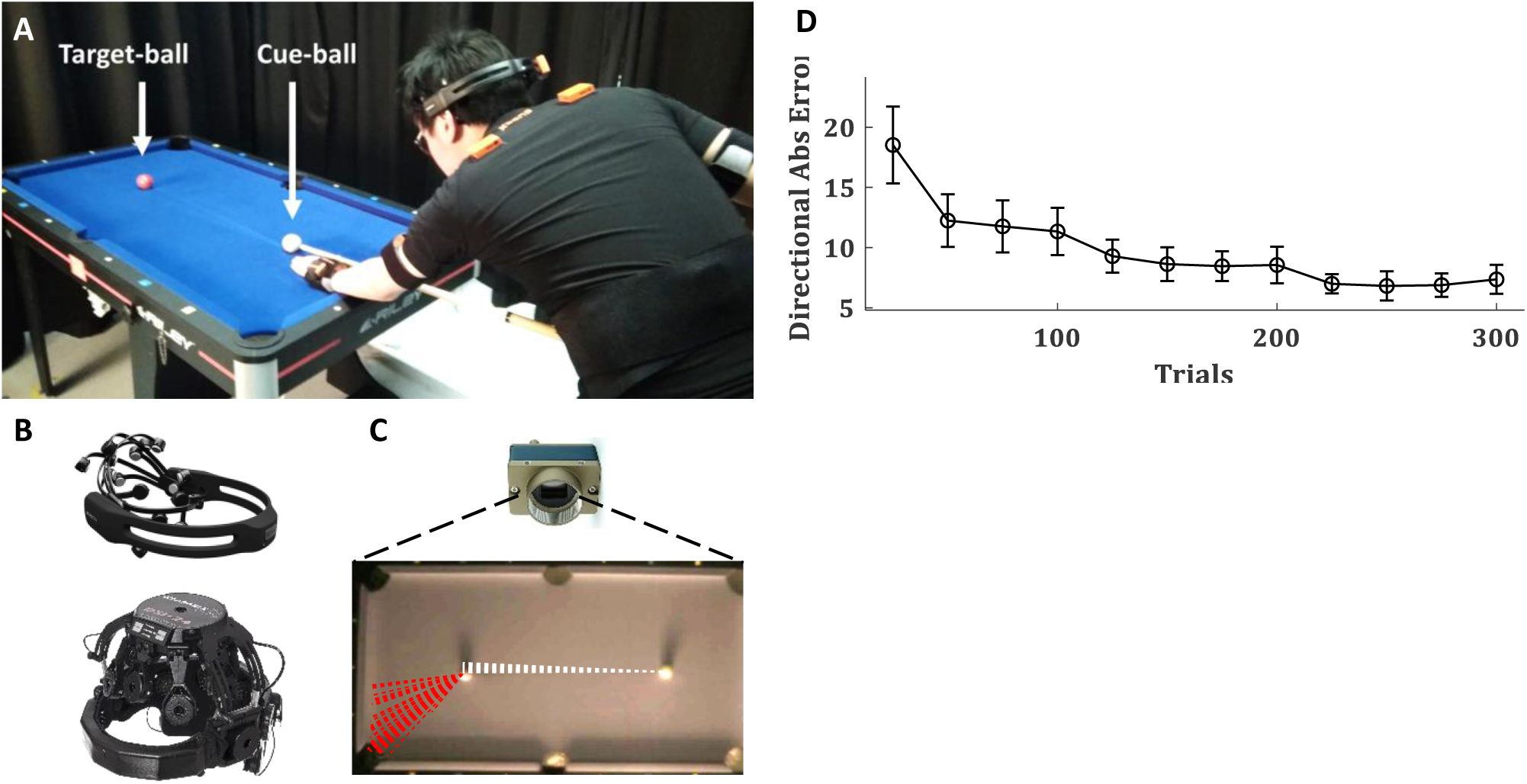
Experimental setup and task performance. (**A**) 30 right-handed healthy subjects performed 300 repeated trials of billiards shoots of the target (red) ball towards the far-left corner. (**B**) Brain activity was recorded with wireless EEG systems: 20 subjects with eMotiv EPOC+ (left) and 10 subjects with Wearable Sensing DSI-24 (right). (**C**) The pool balls were tracked with a high-speed camera. Dashed lines show the trajectories of the cue (white) and target (red) balls over 50 trials of an example subject. (**D**) The mean absolute directional error of the target-ball (relative to the direction from its origin to the centre of the target pocket) over blocks of 25 trials, averaged across all subjects, error bars represent SEM across subjects.

### Balls tracking

The balls movement on the pool table were tracked with a computer vision system mounted from the ceiling. The computer vision camera was a Genie Nano C1280 Color Camera (Teledyne Dalsa, Waterloo, Canada), colour images were recorded with a resolution of 752×444 pixels and a frequency of 200Hz. This Ethernet-based camera was controlled via the Common Vision Blox Management Console (Stemmer Imaging, Puchheim, Germany) and image videos recorded with our custom software written in C++ based on a template provided by Stemmer Imaging. Our software captured the high-performance event timer, the camera frames and converted the images from the camera’s proprietary CVB format to the open-source OpenCV (https://opencv.org/) image format for further processing in OpenCV. The video frames were stored as an uncompressed AVI file to preserve the mapping between pixel changes and timings and the computer’s real-time clock time-stamps were recorded to a text file. Each trial was subject-paced, so the experimenter observed the subject and hit the spacebar key as an additional trigger event to the time-stamps text file. This timing data was later used to assist segmentation of the continuous data stream into trials. The positions of the two pool balls (white cue ball and red target ball) were calculated from the video recordings offline using custom software written in C++ using OpenCV. Then, with custom software written in MATLAB (R2017a, The MathWorks, Inc., MA, USA), we segmented the ball tracking data and extracted the trajectory of the balls in each trial. For each trial, a 20 x 20 pixels (approx 40 x 40 mm) bounding box was set around the centre of the 48 mm diameter cue ball. The time the centre of the ball left the bounding box was recorded as the beginning of the cue ball movement. The pixel resolution and frame rate were thus sufficient to detect movement onset, acceleration and deceleration of the pool balls. The target (red) ball initial position and its position in the point of its peak velocity were used to calculate the ball movement angle (relative to a perfectly straight line between the white cue ball and the red target ball). We subtracted this angle from the centre of the pocket angle (the angle the target ball initial position and the centre of the pocket relative to the same straight line between the balls) to calculate the directional error for each shot.

### EEG acquisition and preprocessing

For the first group of 20 subjects, EEG was recorded at 256Hz using a wireless 14 channel EEG system (Emotiv EPOC+, Emotiv Inc., CA, USA) as we wanted to demonstrate the feasibility of using a consumer-grade system for free behaviour research. In order to then validate our results with a research-grade EEG system, we ran another group of 10 subjects the DSI-24 (Wearable Sensing Inc., CA, USA). With this wireless 21 channel EEG system, EEG was recorded at 300Hz and downsampled to 256Hz to be analysed with the same pipeline as the first group. Since there was no difference in the outcomes between the different systems (see results), we analysed them as a single group, except the system comparison analysis. EEG signals were preprocessed in EEGLAB (https://sccn.ucsd.edu/eeglab; Delorme and Makeig, 2004). EEG signals were first band-pass filtered at 5-35 Hz using a basic FIR filter, and then decomposed into independent component (IC) and artefact ICs were removed with ADJUST, an EEGLAB plug-in for automatic artefact detection (Mognon et al., 2011). Following previous PMBR studies in motor learning (Tan et al., 2014a, 2016; Torrecillos et al., 2015; Alayrangues et al., 2019), all further analysis was performed on the EEG activity over the motor cortex contralateral to the moving arm. As all subjects were right-handed and the movement during the trial was done almost exclusively by the right arm (Haar et al., 2019), we focused on the left motor cortex. Following (Alayrangues et al., 2019) we manually selected for each subject an IC based on its topographies. In order to validate it with a less subjective approach, we repeated the analysis using a single channel, C3 according to the international 10–20 EEG system, which sits over the left motor cortex. For the subjects recorded with the Emotiv system C3 channel was interpolated from the recorded channels with spherical splines using EEGLAB ‘eeg_interp’ function. The two approaches yield the same results, thus, the data reported here is that of the latter. We repeated the analysis over the right motor cortex (ipsilateral to the moving arm contralateral to the stabilizing arm) using C4 according to the international 10–20 EEG system. This analysis yields similar results and is reported in the Supplementary Materials.

### EEG time-frequency analysis

Each block was transformed in the time-frequency domain by convolution with the complex Morlet wavelets in 1 Hz steps. Event-related EEG power change was subsequently calculated as the percentage change by log-transforming the raw power data and then normalizing relative to the average power calculated over the block, as no clear baseline could be defined during the task (Tan et al., 2014a, 2016; Torrecillos et al., 2015; Alayrangues et al., 2019), and then subtracting one from the normalized value and multiplying by 100. While this normalization procedure might be less common than one based on motion-free pre-movement baseline period, it was used by most of the PMBR motor learning studies mentioned above and enabled the natural free-behaviour aspect of the task of self-paced movement, with as many preparatory movements as the subject needs for each shoot, and no go-cues or hold-cues. Event-related power changes in the Beta band (13–30 Hz) were investigated. Since there was no go cue and the subject shot when they wanted, the best-defined time point during a trial was the beginning of the cue ball movement, defined by exiting its bounding box (see *Balls tracking* above). Thus, we used the ball movement onset to estimate movement offset (which could last few hundred milliseconds more due to follow through movement) and looked in the following 2 seconds window for the peak Beta power which should follow the movement termination. The post-movement Beta rebound (PMBR) was defined as the average normalized power over a 200ms window centred on the peak of the power after movement termination (Tan et al., 2016). The PMBR was calculated for each trial before averaging over blocks for further analysis. The time-frequency analysis was performed with custom software written in MATLAB (R2017a, The MathWorks, Inc., MA, USA).

### Multiple groups analysis

To assessed if there may be multiple groups of subjects with different PMBR trends we used generative Bayesian modelling to determine in a data-driven way the structure of the data. We fitted the data with a Gaussian mixture models of one to five components, allowing us to understand if 1,2,3,4 or 5 distinct groups appeared in the distribution or not. To select between these 5 models of different complexity we used two information criteria, the Akaike information criterion (AIC) and its corrected version for small sample size (AICc). AIC estimates the amount of information that is lost while fitting a model and thus can measure the quality of different models relative to each other. In addition to the Bayesian framework, we validate this grouping with unsupervised fuzzy c-means (FCM) clustering, tested for two to ten clusters. FCM assigns a friendship to each data point in each cluster according to its distance from the cluster’s centre, and on iterative process recalculate the clusters’ centres and the friendship until it converges. After convergence, each point is classified into the cluster with which it had the highest friendship. Following Haar et al. (2015), we used the cluster validity index proposed by Zhang et al. (2008). This index uses a ratio between a variation within each cluster and a separation between the fuzzy clusters. The smaller the ratio, the better the clustering.

### Behavioural measures of Motor Skill Learning

We calculated and analysed three know matrices for motor skill learning: movement complexity, lag-1 autocorrelation, and intertrial variability. Movement complexity was defined as the number of degrees of freedom used by the subject as their body move while making the pool shot. For that, we used the manipulative complexity (Belić and Faisal, 2015) over the full-body kinematics. For the analysis of full-body kinematics and its complexity measurements during this task see Haar et al. (2019). Briefly, we applied Principal component analysis (PCA) over the velocity profiles of all body joints and asked how many PCs are needed to explain the variance. The manipulative complexity quantify complexity for a given number of PCs on a fixed scale (C = 1 implies that all PCs contribute equally, and C = 0 if one PC explains all data variability). Lag-1 autocorrelation (ACF(1)) is a lagged Pearson correlation between a signal to itself. In our case, the signal is the directional error of the target-ball relative to the pocket in each trial. Since the estimation of autocorrelations from short time series is fundamentally biased (Kendall, 1954; Marriott and Pope, 1954; van Beers, 2009), we calculated the ACF(1) over the first and the second halves of the learning session (sets of 150 trials, blocks 1-6 and 7-12, respectively) and not in each block of 25 trials. Intertrial variability was defined for each block by the standard deviation over the directional error of the target-ball in all block’s trials. The decay in the intertrial variability was measured from the first block (trials 1-25) to the learning plateau (trials 201-300).

## Results

30 right-handed volunteers, with little to no previous experience playing billiards, performed 300 repeated trials (6 sets of 50 trials each with short breaks in-between) where the cue ball and target ball were placed in the same locations, and subjects were asked to shoot the target ball towards the far-left corner pocket (Figure 1A). During the entire learning process, we recorded the subjects’ brain activity with wireless EEG (Figure 1B), and the balls on the pool table were tracked with a high-speed camera to assess the outcome of each trial (Figure 1C). We divided the trials into blocks of 25 trials (each experimental set of 50 trials was divided into two blocks to increase the resolution in time). The learning curve showed decay in the directional error of the target ball (relative to the direction from its origin to the centre of the target pocket) over trials (Figure 1D).

The PMBR, a transient increase in Beta oscillations over the motor cortex after the end of the movement, was evident in the data (Figure 2A). On average across subjects, there was no clear trend of PMBR (increase or decrease) over learning (Figure 2B). With a data-driven approach, we assessed if there may be multiple groups with different PMBR trends that averaging blends away. We used generative Bayesian modelling to determine in a data-driven way the structure of the data. We fitted to the PMBR data (a 12-dimensional matrix, one data point per run for each subject) a Gaussian mixture models of one to five components and used AIC and AICc to select between these 5 models (see methods). Both information criteria showed that the data followed a bimodal distribution (Figure 2C).

**Figure 2.**
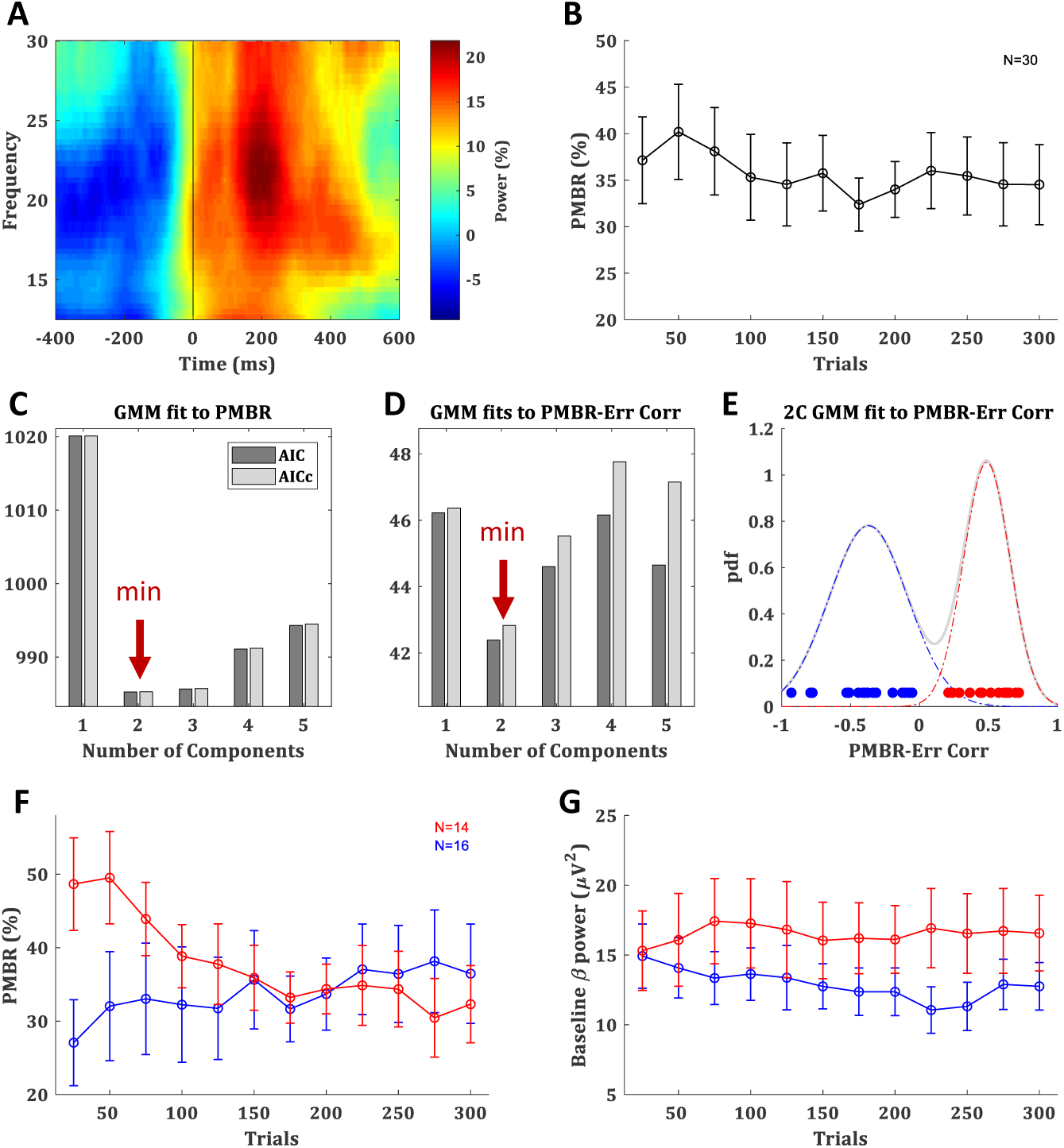
Post-movement beta rebound. (**A**) Time-frequency map of a typical subject aligned to movement offset (ball movement onset), obtained by averaging the normalized power over electrode C3. (**B**) PMBR over blocks (of 25 trials), averaged across all subjects, error bars represent SEM. (**C**) The information criterions (AIC & AICc) of Gaussian mixture model (GMM) fits with 1 to 5 components to the PMBR data. (**D**) The information criterions of GMM fits to the PMBR-Error correlations (**E**) The distribution of subject-by-subject PMBR-Error correlations fitted with two-component GMM (pdf: probability density function). Subjects are color coded based on the two-component model: subjects with negative correlations are in blue (*PMBR Increasers*) and subjects with positive correlations are in red (*PMBR Decreasers*). The grouping was also validated by unsupervised clustering (see main text). (**F**,**G**) PMBR (F) and Baseline beta power (G) of the *PMBR Increasers* (blue) and *PMBR Decreasers* (red) over blocks, averaged across all subjects in each groups, error bars represent SEM.

The most meaningful measure for learning is the PMBR correlation with the performance error, as it accounts for the dependency between this brain signal and the behaviour, and it was reported to show negative correlations in classic adaptation task consistently across individuals (e.g. Tan et al., 2016). The subject-by-subject correlation over blocks between the PMBR and the directional error showed a clear bimodal grouping. While 16 of the 30 subjects showed negative PMBR-Error correlations (as reported in adaptation studies), the other 14 subjects showed positive correlations. Again, we used generative Bayesian modelling to determine the structure of the data. We fitted to the distribution of the PMBR-Error correlations a Gaussian mixture models of one to five components, the information criteria (AIC & AICc) showed that the data followed a clear bimodal distribution (Figure 2D). Each mode corresponded to a grouping of subjects with either all positive and all negative correlation coefficients (Figure 2E). We note that the opposite signs of the correlations reflect opposite dynamics, further justifying a grouping into two distinct groups. This validated our findings with the purely data-driven approach on the multidimensional PMBR data. Since errors decay over learning, the PMBR-Error correlation was negatively correlated with the PMBR dynamic (increase/decrease). Thus, the first group showed a clear trend of PMBR increase over learning (linear model fit: F-statistic vs. constant model = 24 p=0.0006), while the second group showed a clear trend of PMBR decrease over learning (F vs. constant model = 45.1 p=0.00005) (Figure 2F). This was validated with a mixed-design ANOVA model with a between-subjects factor of the group effect, a within-subjects repeated measures factor of the change over blocks, and their interaction. The model yielded a significant interaction (F(11)=6.746 p= 3e-10), but no significance for the between- and within-subjects factors (F(1)=0.27 p=0.61 and F(11)= 1.767 p=0.06, respectively). Thus, for simplicity, we named the groups *PMBR Increasers* and *PMBR Decreasers*.

While we pursued a probabilistic analysis Bayesian framework of data science, to further validate this grouping we also tried a completely different method. We used unsupervised fuzzy c-means (FCM) clustering, tested for two to ten clusters using a cluster validity index based on the ratio between-within cluster variation and between clusters separation (Zhang et al., 2008). The validity index strictly suggested two clusters in the data, which were the same groups found by the Gaussian mixture model: the subjects with the positive and the negative PMBR-Error correlation coefficients. Additionally, since we calculated Beta-power changes as per cent signal change relative to the average power over the block (see methods), the observed group differences might be driven by differences in their baselines. However, we found that this was not the case: there was no real difference in the Beta-power baseline between the groups, in terms of their values and trend over learning (Figure 2G). There was no significant difference between the groups’ Beta-power baseline in any of the blocks (t-test p>0.076) and not in the change of the Beta-power baseline between blocks (t-test p>0.58). This was also validated with a mixed-design ANOVA model which yielded no significant group effect (F(1)=1.286 p=0.27), change over blocks effect (F(11)=0.685 p=0.75) or interaction (F(11)= 1.169 p=0.31). Lastly, we ensured that these groupings were evident with both EEG systems used in the study. The brain activity of 20 subjects was recorded with EPOC+ while the other 10 were recorded with DSI-24 (see methods). From the subjects recorded with the EPOC+ system, 10 subjects were *PMBR Increasers* and the other 10 were *PMBR Decreasers*. From the subjects recorded with the DSI-24 there were 6 *PMBR Increasers* and 4 *PMBR Decreasers*. Correspondingly, there was no correlation between the system and the PMBR-Error correlation (Spearman rank correlation r=0.01 p=0.97).

Based on the EEG data, which suggests two groups of subjects with different PMBR dynamics, we looked for behavioural signatures in the task performance of different learning between these groups. In the task performance metric – the target ball directional error – we found no significant difference between the groups. After learning plateaus, the *PMBR Decreasers* seems slightly more accurate (Figure 3A) and less variable (Figure 3B), though not significantly. Mixed-design ANOVA model yielded no significant group effect (F(1)=0.001 p=0.97) or interaction (F(11)= 0.75 p=0.69) for the absolute directional error. *PMBR Decreasers* seemed to modify their variability (actively control of the exploration-exploitation trade-off, explicitly or implicitly) to improve learning, as evidenced by their high variability in the first block and the very steep decrease towards the second (Figure 3B). Yet, the Mixed-design ANOVA model of the directional variability yielded no significant group effect (F(1)=0.25 p=0.62) or interaction (F(11)= 1.57 p=0.11). The dynamical control of the variability also evident in the trial-to-trial directional changes, where the *PMBR Decreasers* showed much bigger changes over the first 4 blocks (100 trials), therefore using more exploration than the *PMBR Increasers* who made smaller changes from one trial to the next (Figure 3C). Here the Mixed-design ANOVA model yielded close to significance interaction (F(11)= 1.76 p=0.06), and a t-test over the trial-to-trial directional changes in the initial 4 block showed significant group effect (p=0.04).

**Figure 3.**
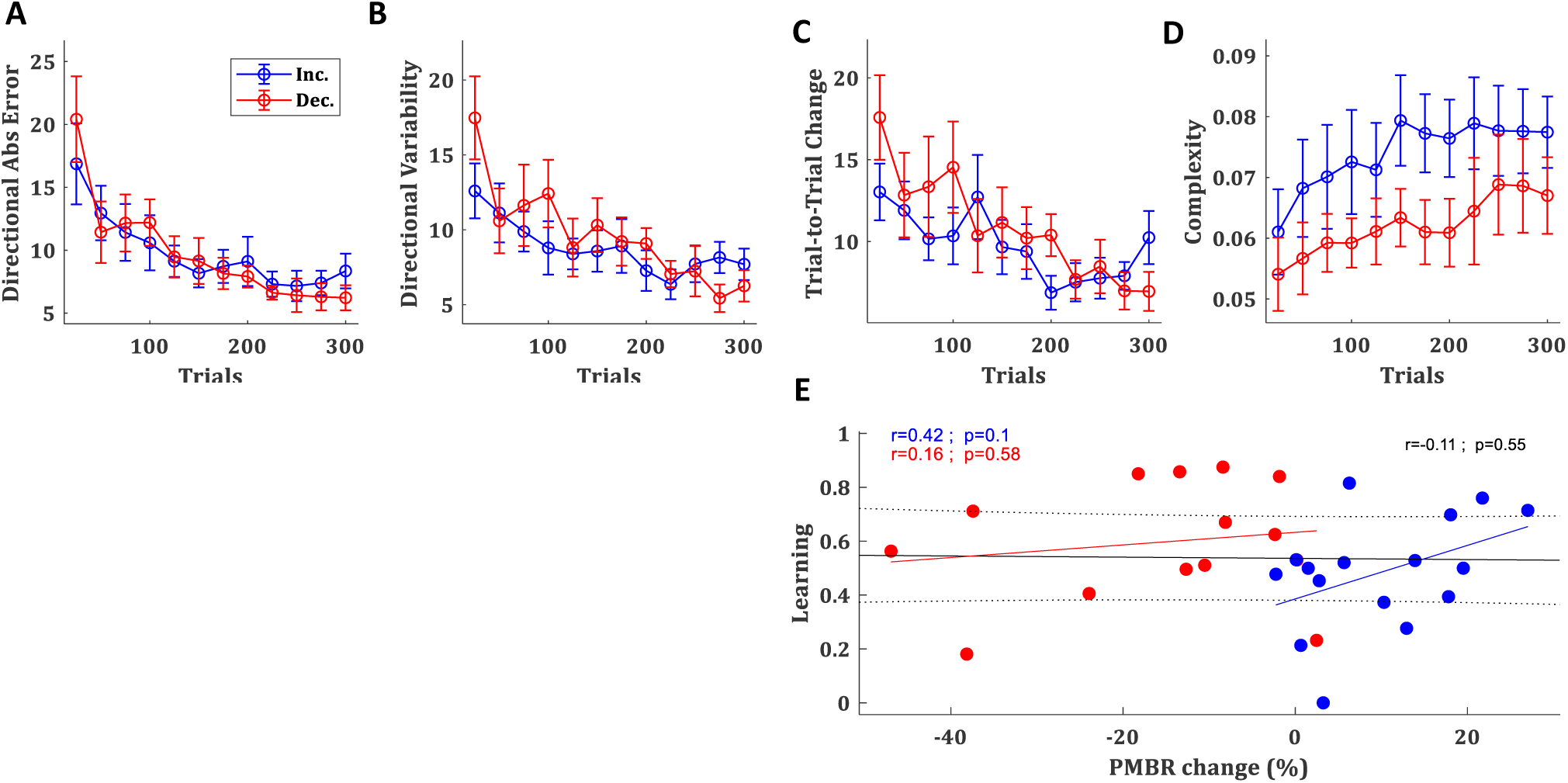
Behavioural differences between the groups. (**A-D**) Directional absolute error (**A**), directional variability (**B**), trial-to-trial directional change (**C**), and manipulative complexity (**D**) of the *PMBR Increasers* (blue) and *PMBR Decreasers* (red) over blocks of 25 trials, averaged across all subjects in each group, error bars represent SEM. (**E**) Correlations between the PMBR change (from the first block (trials 1-25) to the learning plateau (trials 201-300)) and the learning, across all subjects (black line) and within each group.

Learning in the task was defined as the difference between the initial error (over the first block: trials 1-25) and the final error (over the learning plateau: trials 201-300) normalised by the initial error. *PMBR Decreasers* were on average better learners (mean learning rates were 0.48 and 0.6 for the *PMBR Increasers* and *PMBR Decreasers* respectively) though the group difference was not significant (t-test p=0.17). We explored the correlation between learning and the PMBR change over blocks (the difference between the final PMBR over the learning plateau: trials 201-300, and the initial PMBR over the first block: trials 1-25). Across all subjects, we found no correlation between the learning rate and the PMBR change (r=-0.11 p=0.55). When considering each group separately, for the *PMBR Decreasers* there was no clear trend (r=0.16 p=0.58), but the *PMBR Increasers* showed a clear trend (though non-significant) of positive correlation of the PMBR change with learning (r=0.42 p=0.1, Figure 3E). This means that within the *PMBR Increasers* group subjects who had a higher PMBR increase also showed more learning.

Next, we set to study metrics of skill-learning which might suggest differences in the learning mechanism between the groups. First, we tested the complexity of the movement – i.e. the number of degrees of freedom used by the subject – since the use of multiple degrees of freedom in the movement is a hallmark of skill learning (Bernstein, 1967). For that we used the manipulative complexity (Belić and Faisal, 2015) over the full-body kinematics (see Haar et al. (2019) for the analysis of full-body kinematics and its complexity measurements during this task). While the manipulative complexity is increasing with learning for all subjects, *PMBR Increasers* tended to have higher complexity in their movement, i.e. use more DoF, throughout the training session (t-test p<0.05, Figure 3D).

Second, we explored the lag-1 autocorrelation (ACF(1)) of the performance measure (in our case, the directional error of the target-ball relative to the pocket) which was suggested as an index of skill, where close to zero values corresponds to high skill (van Beers et al., 2013). The logic behind this measure is that as skill evolve subjects are less susceptive to noise from the previous movement. We calculated the ACF(1) over the first and the second halves of the learning session (sets of 150 trials, blocks 1-6 and 7-12, respectively). The ACF(1) values of both groups were significantly greater than zero during both halves of the session (t-test p<0.01), as expected for naïve participants (Figure 4A). The initial ACF(1) values of the *PMBR Decreasers* were higher than those of the *PMBR Increasers*, though not significantly (t-test p=0.06). But, the decay in the ACF(1) from the first half of the training session to the second was significantly higher for the *PMBR Decreasers* (t-test p<0.01, Figure 4B). This was also validated with a mixed-design ANOVA model which yielded no significant overall group effect (F(1)=0.119 p=0.73), but a significant change over the two halves (F(1)=7.79 p=0.009) and a significant interaction (F(1)= 8.393 p=0.007).

**Figure 4.**
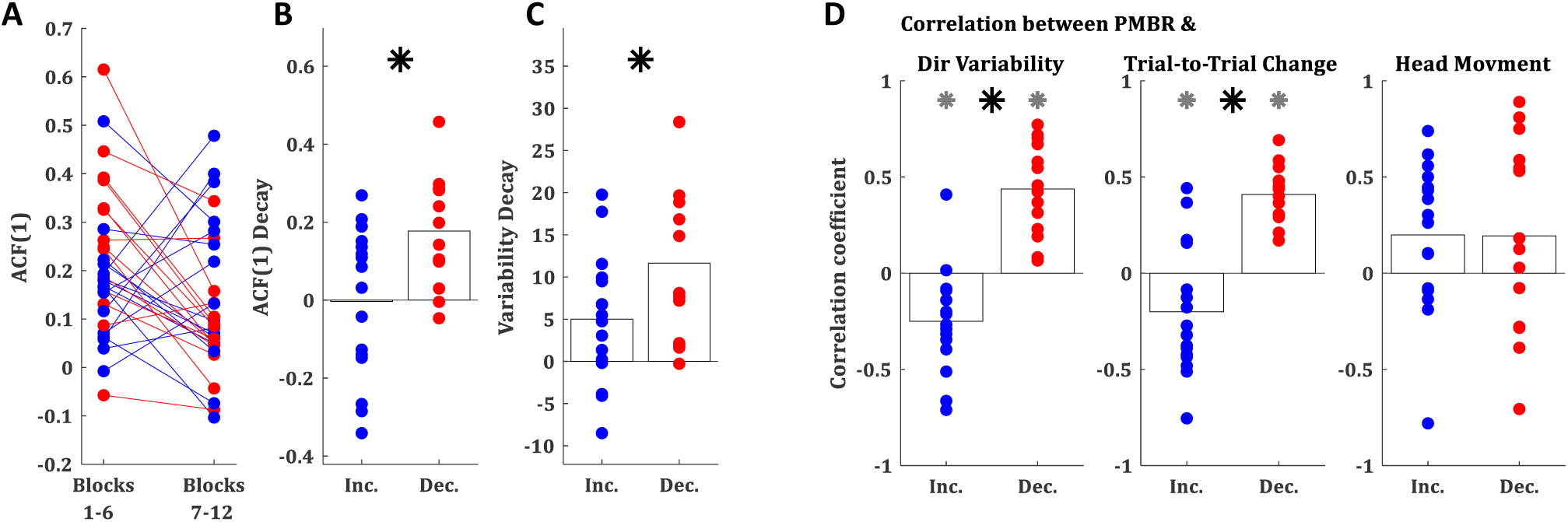
Behavioural differences between the groups. (**A**) Lag-1 autocorrelation of the target ball direction over the first and the second half of the training session (blue: *PMBR Increasers*; red: *PMBR Decreasers*). (**B**) Decay of the lag-1 autocorrelation from the first to the second half of the training session. (**C**) Directional variability decay from the first block (trials 1-25) to the learning plateau (trials 201-300). (**D**) Correlation coefficients over blocks for all individual subjects between the PMBR and the directional variability (left), trial-to-trial directional change (middle), and head movements (right). Grey asterisk indicates group correlations significantly different than zero. Black asterisk indicates significant difference in the correlation coefficients between the groups.

A third behavioural measure which can differentiate between learning mechanisms is the decay in the intertrial variability over learning, which is a known feature of skill learning (Deutsch and Newell, 2004; Müller and Sternad, 2004; Cohen and Sternad, 2009; Guo and Raymond, 2010; Shmuelof et al., 2012; Huber et al., 2016; Sternad, 2018; Krakauer et al., 2019). The decay in the intertrial variability (measured from the first block (trials 1-25) to the learning plateau (trials 201-300)) was also significantly larger in the *PMBR Decreasers* (t-test p<0.05, Figure 4C). We further tested the link between the intertrial variability structure and the reported grouping by correlating the block-by-block directional variability and PMBR values within subjects. The *PMBR Increasers* showed negative correlations over blocks between the PMBR and the directional variability, while the *PMBR Decreasers* showed positive correlations, leading to a very significant difference between the groups (t-test p<0.0001, Figure 4D left). The same trend was evident for the trial-to-trial directional changes (t-test p<0.0001, Figure 4D middle). We also used the same correlation approach to control for head movements contamination of the PMBR dynamics. We looked for correlation over blocks between the PMBR and the peak head acceleration during the same time interval. Here we found no significant correlations for either of the groups (Figure 4D right), and most importantly, no difference between the groups (t-test p=0.99).

## Discussion

In this paper, we detected brain activity signatures for motor learning in the complex real-world task of playing pool billiards. Our results produce new insights into motor learning by revealing two types of motor learners with different EEG dynamics in their PMBR over learning: *PMBR Increasers* and *PMBR Decreasers*. These groups were defined by their PMBR dynamic, and the grouping was validated over the correlation between the dynamics of their PMBR and their performance errors. While the groups showed no difference in the overall task performance – as measured by the directional errors of the ball – there were clear task-level differences between the groups in measures of skill learning which suggest differences in the underlying learning mechanisms.

The two known main mechanisms that drive motor learning – error-based learning and reward-based reinforcement learning – are engaging different neural processes (Doyon et al., 2003; e.g. Doyon and Benali, 2005; Uehara et al., 2018). While both mechanisms can contribute to learning in any given task, controlled laboratory-tasks are usually designed to induce the predominance of one mechanism over the other. In motor adaptation tasks the dominant mechanism is error-based learning, guided by an internal forward model which is updated based on sensory-prediction errors; while in tasks often addressed as skill-learning (such as sequence-learning, curve-tracking, and force-tracking) the dominant mechanism is reward-based learning where the controller learns form reinforcement of successful actions (Krakauer and Mazzoni, 2011; Haith and Krakauer, 2013). PMBR was reported to increase over learning in adaptation error-based learning tasks (e.g. Tan et al., 2014a, 2016; Torrecillos et al., 2015), showing negative correlations with the decreasing errors. On the other hand, in skill-learning tasks it was reported to decrease (itself or its magnetic resonance spectroscopy correlate) over the learning (e.g. Floyer-Lea et al., 2006; Kranczioch et al., 2008; Kolasinski et al., 2019). PMBR is positively correlated with magnetic resonance spectroscopy-measured GABA concentration (Gaetz et al., 2011; Cheng et al., 2017). This may be due to the general correlation of Beta activity with GABAergic activity (Roopun et al., 2006; Yamawaki et al., 2008; Hall et al., 2010, 2011). We raise the possibility of a more nuanced link of GABA to motor learning: namely that the two diverging PMBR dynamics (increase vs. decrease) reflect that GABA activity is a distinguishing feature of different motor learning mechanisms. These may be reflections of GABAergic projections from different subcortical regions, cerebellum for error-based adaptation and basal ganglia for reward-based reinforcement learning (Doyon et al., 2003; Doyon and Benali, 2005).

Here, we found PMBR dynamic differences between groups of subjects performing the same task and explored it as a potential signature of motor learning mechanisms. In the data recorded during real-world motor learning in the current study, we found two groups of subjects: *PMBR Increasers* and *PMBR Decreasers*. The *PMBR Increasers* had low initial PMBR amplitudes and showed an increase over learning negatively correlated with the decreasing directional errors (r=-0.40±0.26). Following previous PMBR literature reported above, we presumed that these subjects used error-based adaptation as their dominant learning mechanism. The *PMBR Decreasers* had higher initial PMBR amplitudes and showed a decrease over learning positively correlated with the decreasing directional errors (r=0.47±0.17). Again, following previous PMBR literature, we presume that these subjects used reward-based learning as their dominant learning mechanism. As this mapping is highly speculative, we further explored the performance of the different groups in the task, looking for signatures of differences in the learning mechanisms in use. While the main text results are based on the PMBR over the left motor cortex (contralateral to the moving arm), we repeated the analysis over the right motor cortex (ipsilateral to the moving arm contralateral to the stabilizing arm). In line with previous literature showing similar PMBR trends between the two hemispheres (e.g., Jurkiewicz et al., 2006; Gaetz et al., 2010), the results over the right motor cortex (reported in the Supplementary Materials) replicated those of the left motor cortex, straightening the robustness of the different PMBR trends.

While there were no significant differences between the groups in their initial errors or their total learning, there was a clear group difference in the learning process. These behavioural differences can support the notion of differences in the predominant learning mechanism. First, we looked for group differences in the number of degrees of freedom of the body movement used while making the pool shot. Since the pioneering work of Nikolai Bernstein, who found that professional blacksmiths use high variability in their joint angles across repetitive trials to achieve low variability in their hammer trajectory endpoint, it is known that as skill evolves one learns to use more degrees of freedom in the movement (Bernstein, 1967). Using the full-body kinematics in this task from our previous study (Haar et al., 2019), we found that while over learning both groups learned to use more degrees of freedom in their movement, throughout the learning session there was a clear group difference where the *PMBR Increasers* used more degrees of freedom in their movement (Figure 3D).

We used the lag-1 autocorrelation (ACF(1)) as a second biomarker for the difference in the learning prosses between the groups. ACF(1) was suggested as an index of skill learning, measuring the optimality of trial-by-trial motor planning (van Beers et al., 2013). ACF(1) of zero indicates optimal performance. What ACF(1) measures is the correlation between the errors in consecutive trials, and thus could be a good metric to dissociate between error-based adaptation (where we gradually decrease the error from one trial to the next) to reinforcement learning (where an error should lead to exploration). As expected for naïve participants, the ACF(1) values of both groups during both halves of the session were significantly greater than zero (Figure 4A). More importantly, while the *PMBR Increasers* showed no significant difference in the ACF(1) between the two halves of the session, the *PMBR Decreasers* showed a significant decay (Figure 4B). This decays difference is a behavioural indication for learning mechanism differences between the groups.

Third, the intertrial variability patterns were in line with the suggestion of different learning mechanisms. A decay in the intertrial variability a known feature of skill learning (Deutsch and Newell, 2004; Müller and Sternad, 2004; Cohen and Sternad, 2009; Guo and Raymond, 2010; Shmuelof et al., 2012; Huber et al., 2016; Sternad, 2018; Krakauer et al., 2019), but not of adaptation. Here, the *PMBR Decreasers* (presumably reward-based learners) showed more decay in their intertrial variability over learning (Figure 3B & Figure 4C). Additionally, the trial-to-trial directional changes over the first 4 blocks (100 trials) were much higher for the *PMBR Decreasers* than the *PMBR Increasers* group, suggesting that the first group used more exploration while the second made smaller changes from trial-to-trial (Figure 4C). This latter behaviour would be expected when learning predominantly by error-based adaptation.

Laboratory-tasks are usually designed to look or characterise a specific learning mechanism (which is being studied) for all subjects, using different types of feedback and perturbation manipulations (Huang et al., 2011; e.g. Galea et al., 2015; Kim et al., 2019). In contrast, the way we started to study real-world motor learning here, which mechanisms are used and to what extent is unknown a priori. However, we know that they probably involve multiple high- and low-level learning mechanisms (Krakauer and Mazzoni, 2011; Haith and Krakauer, 2013), where different subjects might emphasize one learning modality over the other.

In our pool playing paradigm, subjects could have performed error-based adaptation as they learned from the directional error of the target ball in each trial, but they also could have performed reward-based learning as they learned a novel control policy to use the cue and their body joints while making a shot by reinforcement of successful actions. In the following, we will discuss how we could map the distinct groups of learners we discovered in our real-world task into the above learning frameworks (error-based and reward-based). We speculate that the group that showed the neural patterns which were previously reported in error-based motor adaptation (PMBR increase, (Tan et al., 2014a, 2016)) and behavioural patterns of error-based adaptation (e.g. no decay in AFC, small decay in intertrial-variability, low trial-to-trial change) – probably used more error-based adaptation to adapt an existing motor control policy. At the same time, the group that showed neural patterns which were previously reported in reward-based motor skill learning (PMBR decrease, (Kranczioch et al., 2008)) and behavioural patterns of motor skill learning tasks (e.g. decay in AFC, decay in intertrial-variability, high trial-to-trial change) – probably used more reinforcement reward of successful actions for learning a new control policy.

We recently showed in a machine learning study how simultaneous reinforcement learning and error-based learning can efficiently be used to learn to control multi-joint muscle activities to learn to control an arm (Abramova et al., 2012, 2019). That work suggested when adaptation should occur: if a “similar enough” controller to achieve the task is already present (e.g. from other motor learning experiences) the existing controller is adapted then learning should have an error correction signature. In contrast, the absence of a suitable controller for the task either spawned the generation of a new controller or switching between multiple somewhat suitable controllers. We may see similar effects at work in this present human study for the two groups of learners. Thus, our real-world task merit further investigation not only in terms of the neuroscience of learning, but also in light of robot and machine learning algorithms that could explain the combination of these learning paradigms or even an entirely new process.

Recent studies suggest that event-related desynchronizations and synchronizations, such as PMBR, are driven by Beta bursts (Little et al., 2019; Seedat et al., 2020; Wessel, 2020) which carry more information than the trial-averaged band oscillation. At the same time, a recent study suggested spatial differences between Beta oscillations that reflect implicit and explicit learning (Jahani et al., 2020). These recent developments highlight the potential for capturing neural signatures of learning in EEG Beta. To further validate the current findings, future studies will need to compare the PMBR dynamics during learning of the same paradigm with different dominant mechanism, forced by experimental trickery (i.e. using feedback manipulations and constraints) in laboratory-tasks or real-world task in a virtual reality environment, where feedback manipulations can be applied (Haar et al., 2020).

The transition from a highly controlled lab-based task to a more ecological free-behaviour task introduces many challenges which led to a few limitations in the design. First, event-related EEG power changes are ideally normalized relative to a motion-free pre-movement baseline period. Since we were trying to keep the task as ecologically valid as possible, we choose not to force on the subjects a period of quiescence before each shot. Instead, we normalized relative to the average power. This follows a common normalization protocol in studies of PMBR during motor learning in lab-based tasks (Tan et al., 2014a, 2016; Torrecillos et al., 2015; Alayrangues et al., 2019). Since the same normalization was applied to all blocks of all subjects, we believe that the normalization protocol could not have affected the within-subject PMBR trends in a way that would change the results. Second, movement has termination is also not perfectly defined, as subjects could follow through, or not. To address it, we used the ball-movement onset to define the movement offset and defined the PMBR based on the peak of the power in the following two seconds. Thus, even if the follow-through lasted a few hundred milliseconds, the PMBR was well within the window.

Finally, the results of the current study are correlational and cannot, by design, establish a causal role of PMBR in motor learning or motor learning causing PMBR. We propose, however, that going forward that brain stimulation at the Beta band can be used to manipulate the PMBR in order to infer causality (Pogosyan et al., 2009; Tan et al., 2014b; Herrmann et al., 2016). Similarly, differential studies with patient groups with evidence of an impaired Beta activity, such as Parkinson (Heinrichs-Graham et al., 2014), stroke (Rossiter et al., 2014), Autism Spectrum Disorder (Gaetz et al., 2020), or Schizophrenia (Robson et al., 2016), can also provide evidence for evaluating causality. We believe that our natural task approach here will be facilitating working with such patients’ groups instead of using the artificially construed tasks of clinical settings.

## Conclusions

In this mobile brain activity study in a pool playing task, we demonstrate the feasibility and importance of studying human neuroscience in-the-wild, and specifically in naturalistic real-world motor learning. We highlight that real-world motor learning involves different neural dynamics for different subjects, which were previously associated with different learning mechanisms in different tasks. Presumably, the individual subject’s proportion of applying the two learning mechanisms could be revealed by the overall trend of the PMBR over learning. It suggests that real-world motor learning involves multi-modal learning mechanisms which subjects combine in new ways when faced with the complexity of learning in the real-world, and different subjects emphasize one mechanism over the other.

## Acknowledgements

We thank our participants for taking part in the study. We thank Camille M. van Assel and Marlene Gonzalez for their contribution to the data collection. We acknowledge the technical support by Alex Harston and Chaiyawan Auepanwiriyakul. The study was enabled by financial support to a Royal Society-Kohn International Fellowship (NF170650) and by eNHANCE (http://www.enhance-motion.eu) under the European Union’s Horizon 2020 research and innovation programme grant agreement No. 644000. This manuscript has been released as a pre-print at bioRxiv (Haar and Faisal, 2020). Written informed consent was obtained from the individual in Fig1 for the publication of any potentially identifiable images included in this article.

## Supplementary Material

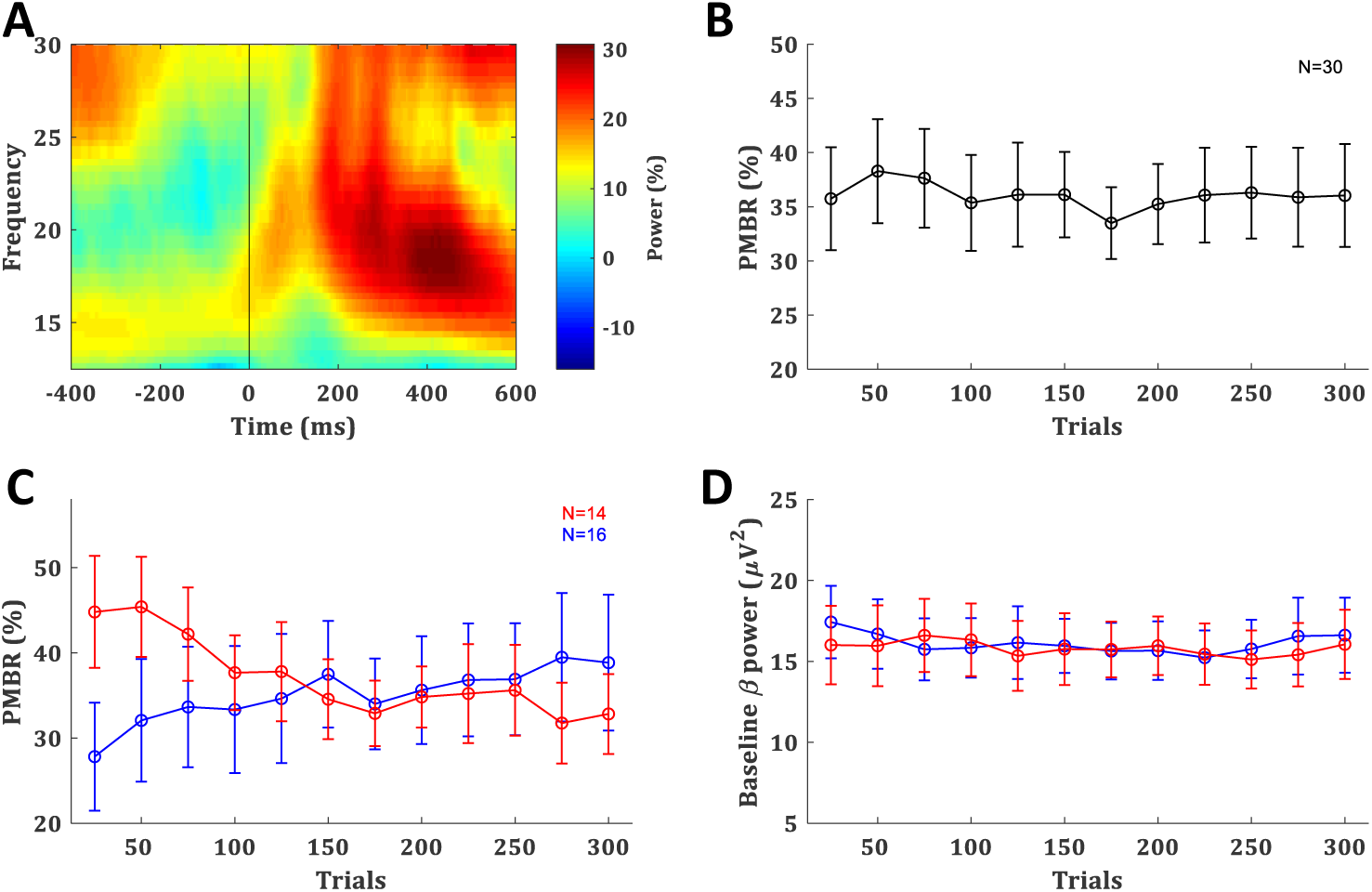
Right Motor Cortex Post-movement beta rebound. (**A**) Time-frequency map of a typical subject aligned to movement offset (ball movement onset), obtained by averaging the normalized power over electrode C4. (**B**) PMBR over blocks (of 25 trials), averaged across all subjects, error bars represent SEM. (**C**,**D**) PMBR (C) and Baseline beta power (D) of the PMBR Increasers (blue) and PMBR Decreasers (red) over blocks, averaged across all subjects in each groups, error bars represent SEM.

